# Converting lateral scanning into axial focusing to speed up 3D microscopy

**DOI:** 10.1101/2020.03.06.981472

**Authors:** Tonmoy Chakraborty, Bo-Jui Chang, Stephan Daetwyler, Etai Sapoznik, Bingying Chen, Kevin M. Dean, Reto Fiolka

## Abstract

In optical microscopy, the slow axial scanning rate of the objective or the sample has traditionally limited the speed of 3D volumetric imaging. Recently, by conjugating either a movable-mirror to the image plane or an electrotuneable lens (ETL) to the back-focal plane respectively, rapid axial scanning has been achieved. However, mechanical actuation of a mirror limits axial scanning rate (usually only 10-100 Hz for piezoelectric or voice coil based actuators), while ETLs introduce spherical and higher order aberrations, thereby preventing high-resolution imaging. Here, we introduce a novel optical design that can transform a lateral-scan motion into a spherical-aberration-free, high-resolution, rapid axial scan. Using a galvanometric mirror, we scan a laser beam laterally in a remote-focusing arm, which is then back-reflected from different heights of a mirror in image space. We characterize the optical performance of this remote focusing technique and use it to accelerate axially swept light-sheet microscopy (ASLM) by one order of magnitude, allowing the quantification of rapid vesicular dynamics in 3D.

## Introduction

The ability to rapidly change the focal plane of an optical imaging system is of great interest to optical microscopy, machine vision and laser machining. Refocusing is traditionally achieved by mechanically translating the objective or the sample. For high-speed imaging, this however becomes rate limiting due to slow dynamics of moving objects of high inertia.

A potential way to alleviate this problem is through remote focusing, which is an emerging technique where optical refocusing is carried out without moving the primary objective or the sample^1,2^. Primarily, this is realized by introducing a wavefront modification either in Fourier space (conjugated to the backfocal plane of the objective) or in image space. The former is typically achieved either by tunable acoustic gradient index of refraction lenses^3,4^, electro-tunable lenses (ETL)^5^ or deformable mirrors^6^ and the latter by moving a mirror axially in a remote image space. Advances in ETL design have led to hundreds of kHz axial scan rates in resonantly operated devices^5,7,8^. However, all ETL designs so far approximate only a quadratic phase function for defocus, which cannot account for the higher order spherical aberrations that need to be addressed in order to maintain a diffraction-limited focus. While deformable mirrors (DM) are capable of producing the more complex wavefronts that are required for aberration free remote focusing, there are trade-offs between speed and actuator stroke: rapid DMs typically have limited stroke in the range of one wavelength^6^, which in turn limits the achievable axial focusing range. On the other hand DMs with large stroke actuators, thus capable of large focus changes, are typically much slower.

Using a carefully pupil matched remote objective, aberration free remote focusing can be achieved by moving a small mirror in the image space of the remote objective. Depending on the actuation technology, scan speeds from 10-1000Hz have been reported, but here also exists a tradeoff between actuation amplitude and scan rate, and currently no mechanical scan technology is in existence that could go beyond the fastest reported rates.

However, especially for the field of neuroscience, three dimensional multiphoton microscopy of membrane voltage activities (with dynamics on the time scale of 1 ms or less)^9^ or cerebral blood flow requires more rapid scan technologies in all three dimensions. This has spurred the development of very fast lateral scan technologies^10–12^, and recently also axial refocusing within ∼5 nanoseconds using a reverberation loop^13^. However technological (pulse repetition rate) and photophysical (fluorescence lifetime) limitations restrict reverberation microscopy to a maximum of ∼10 focal planes. In addition, like acoustic gradient lenses or ETLs, reverberation microscopy employs only a quadratic phase function for re-focusing, and the resulting spherical aberrations limit it to low resolution imaging. Lastly, reverberation microscopy creates a series of differently focused laser spots with an exponentially decaying intensity. As such it is best suited for focal planes that are spaced by one scattering mean free path, typically 100 microns in brain tissue. For smaller degrees of refocusing, very uneven illumination of the different focal planes would result.

Therefore, we still see a need for a high-resolution axial scan technology that has the ability to reach scanning rates comparable to the fastest lateral scan technologies while avoiding spherical aberrations. Since very fast lateral scan technologies have been established and are available (e.g., electro-optic deflectors and mirror galvanometers), we asked ourselves if an optical system could be designed that can transform a lateral scan motion into axial refocusing. The simplest method we could envision is to take the concept of aberration free remote focusing, and instead of moving the corresponding remote mirror axially, we scan a laser spot laterally over a stationary mirror. If the height of the mirror (i.e. the axial distance to the objective lens) varies in the scan direction, the laser spot is defocused, as required for remote refocusing. If on the return path of the back-reflected light the lateral scan component is perfectly compensated, a pure axial scan motion at the rate of the employed (lateral) scanning technology is obtained.

In this manuscript, we describe two variants of this concept, one capable of performing discrete axial steps using a stair-case shaped mirror and one capable of continuous axial scanning using a tilted planar mirror. The former allows in principle arbitrarily large axial step sizes over a finite number of steps, whereas the latter method allows for an arbitrary number and size of axial steps, although over a more limited scan range. We present numerical simulations and experimental measurements of both scan technologies. Experimentally, we explore the version with the tilted planar mirror. Combined with resonant galvanometric scanning, we demonstrate axial focusing at a rate of 12 kHz. We further leverage our new scan mechanism to speed up the framerate of Axially Swept Light Microscopy^14,15^ and use this form of high-resolution light-sheet microscopy to image rapid intracellular dynamics.

## Concept

Figure 1(a) shows a schematic representation of our experimental setup, which consists of a two arms, right-angled to each other, with a 4f telescope and an objective in each. The remote focusing arm contains a galvanometric scanning mirror (GSM) and an air objective lens (OBJ1), while the other arm, which will be referred to as illumination arm, consists of a pupil matched water objective (OBJ2). These two arms are aligned such that the GSM is conjugate to the back focal plane of both objectives. Laser foci with different axial position emerging from OBJ2 are then observed with two imaging systems, one for direct transmission measurements of the foci and one to observe fluorescence in an orthogonal direction. Both imaging systems consist of water immersion objectives, tube lenses, a camera and appropriate optical filters.

**Figure 1.**
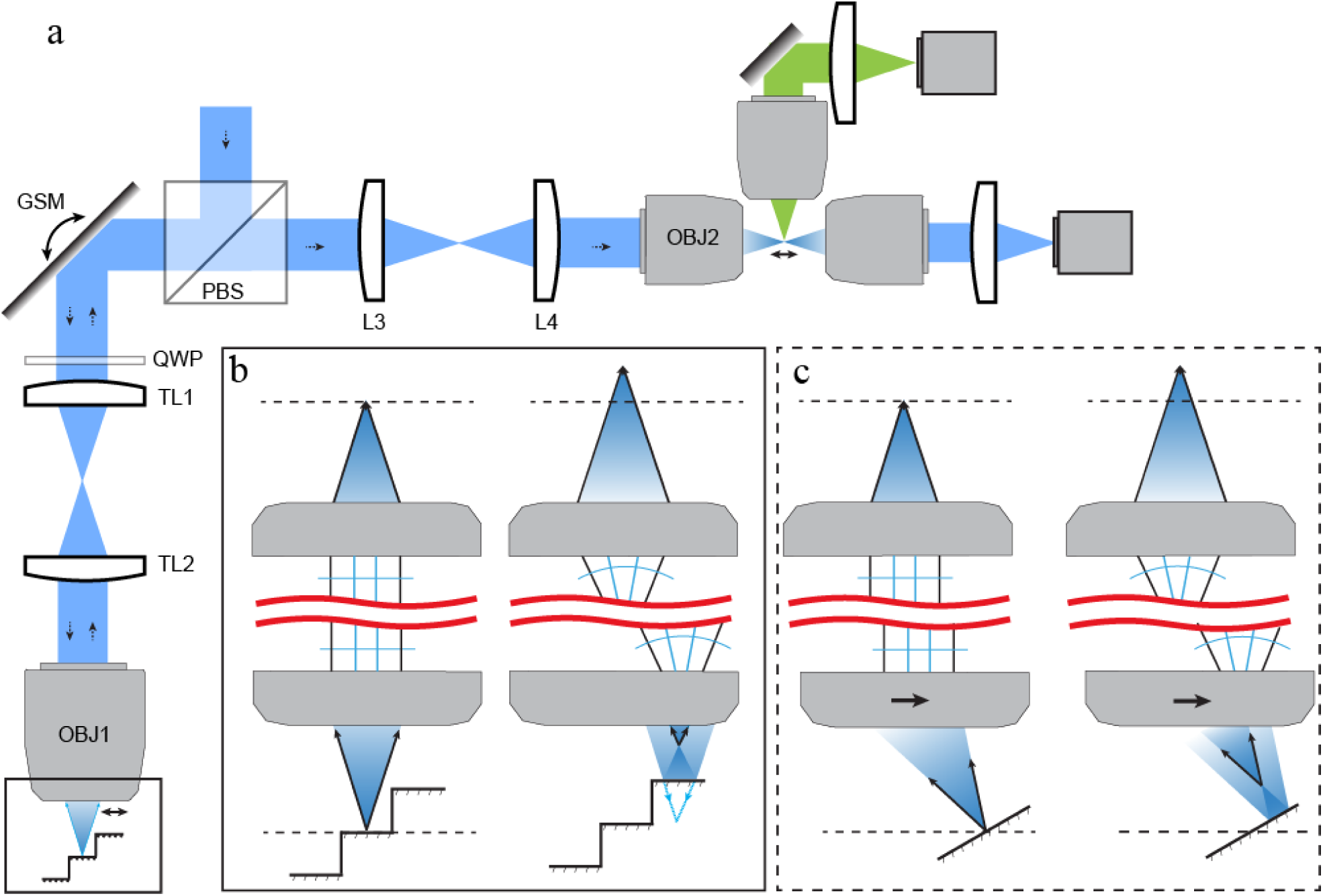
Schematic illustration of remote focusing approach. Setup: (a) A collimated laser beam is delivered into the setup by a beam-splitter (BS) and onto a galvanometric scanning mirror (GSM), which is imaged into the back focal plane of an air objective (OBJ1). Scanning the GSM rasters the focus in one dimension as shown by the double-headed arrow in the boxed front focal space of OBJ1. A step mirror reflects the light with different amounts of defocus back into the objective, which then travels through the 4f system onto the GSM, where it is descanned by, which removes the lateral scan motion and only the axial component remains. The GSM is then again imaged onto the back focal plane of a water dipping objective (OBJ2). As OBJ1 and OBJ2 are pupil matched, OBJ2 forms an aberration free image of the focus (as formed by OBJ1) in the sample space. (b) Zoomed in view of the boxed region from (a). Panel on the left shows the focus of the light at its nominal focus. Black arrows show returning marginal rays after reflection. Each step on the mirror results in a focus spot in the sample plane with a displaced axial position. (c) Alternative configuration with a tilted mirror that allows continuous axial scanning. Here, the remote objective OBJ1 is slightly shifted off the optical axis to create a tilted focus that is incident normal to the mirror surface. Scanning this focus laterally results in a change of focus, as illustrated by the black arrows.

In the setup, an incoming collimated laser beam is reflected by a polarizing beam splitter into the remote-focusing arm where the beam is scanned by the GSM, which in turn produces a lateral scan motion of the laser focus generated by OBJ1. For discrete axial refocusing, a step-mirror is placed in the focal plane of OBJ1 (Figure 1b, left side). When the laser spot is incident on a step that is exactly in the focal plane, the back-reflected laser becomes a collimated beam in infinity space and forms a laser spot in the nominal focal plane of OBJ2. Importantly, as the returning laser beam is de-scanned by the GSM, no lateral scanning motion is imparted on the focus formed by OBJ2 (i.e. the scan motion is purely axial). If the laser beam is scanned to a step that does not coincide with the nominal focal plane, a laser focus away from the focal plane is formed (Figure 1b, right side). The returning laser beam is now converging or diverging in infinity space which in turn causes the laser focus formed by OBJ2 to be axially refocused as well. For an optical setup where the pupil of the two objectives have been accurately matched (in short, the image space from the air objective OBJ1 has to be demagnified by a factor 1.33 into the image space of the water objective OBJ2, according to Botcherby et al.^1,2^), refocusing free of spherical aberrations can be achieved.

For continuous axial refocusing, the step mirror is replaced by a slightly tilted, planar mirror (Figure 1c). The incoming laser focus is also tilted such that it is incident in a direction normal to the mirror surface. In our setup, this tilt is achieved by slightly translating OBJ1 with respect to the incoming laser beam (Suppl. Note 1 and 2). For a beam position where the laser focus is formed on the mirror surface, a collimated beam is formed in infinity space and a laser spot at the nominal focal plane of OBJ2 is formed (Figure 1c, left side). If the beam is scanned laterally, a focus away from the mirror plane is formed (Figure 1c, right side), which in turn causes the beam in infinity space to converge or diverge and leads to an axially displaced focus formed by OBJ2. For small tilt angles, we found numerically (Supplementary Note 1) and experimentally that the focusing remains free of spherical aberrations. For the continuous z-scanning method using a tilted mirror, the angular aperture of OBJ1 has to be larger than that of OBJ2 to permit tilting of the laser focus without giving up any numerical aperture of OBJ2. The achievable axial scan ranges scale approximately as the field of view of OBJ1 times the tangent of the mirror tilt angle. Thus, the technique benefits from objectives and scanning systems that possess a large FOV (Supplementary Note 2). In contrast, using the step mirror, in principle arbitrarily large axial displacements can be realized, with the caveat that for large amounts of defocus, wider steps are needed. This requirement puts practical limits on how many steps can be used for a given field of view of the remote objective (Supplementary Note 3).

## Results

To test the axial focusing technique, we placed a planar mirror that was tilted by 7.5 degree with respect to the image plane, and OBJ1 was carefully laterally translated such that the returning beam retraced the same optical path as in the non-titled case. Scanning the GSM in discrete steps allowed us to produce a series of axially displaced laser spots that span a range of 43 micrometers, as shown in Figure 2a-b. Measuring the full width half maxima of the profiles (364±4.4 nm in x, 371±16 nm in y) confirmed a performance close to the diffraction limit. Lastly, we applied a sawtooth waveform to the GSM, which generated a linear axial line scan (Figure 2c). In the example shown, the beam was scanned to the limit of the field of view of our scan optics, hence field curvature effects became noticeable. As a consequence there was a slight tilting of the foci at the end of the scan range (Supplementary Note 1 and Supplementary Figure 2). The scan range for the discrete axial foci (shown in Figure 2a) is over a range where field curvature of our lateral scanning system is minimal.

**Figure 2:**
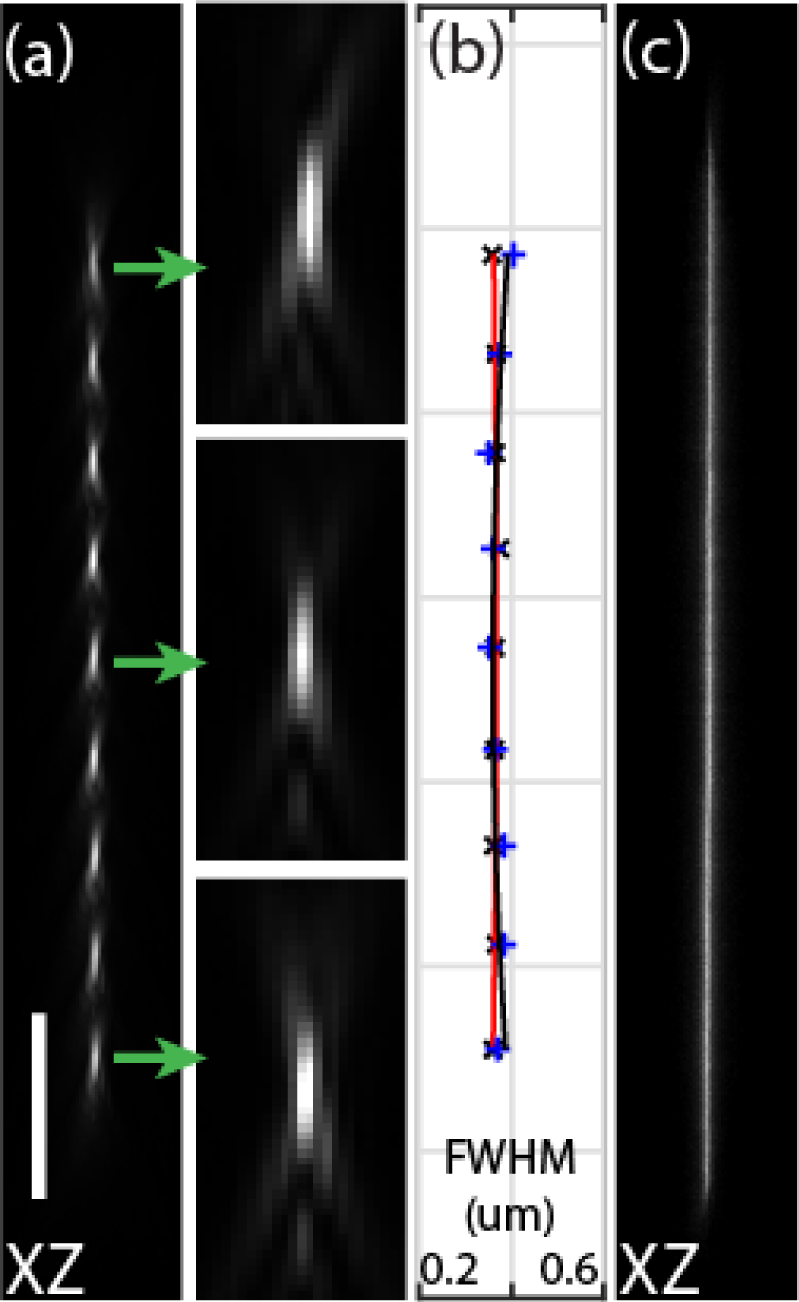
Remote focusing with a planar mirror inclined at 7.5 degrees to the optical axis: (a) PSFs generated form discrete scanning lateral scanning steps. Green arrows depicting the zoomed-in view. (b) Plot showing the full-width-half-max (FWHM) of the PSFs at different axial focus positions. FWHM of the PSF calculated in x (Red) and y (Black) direction. (c) Continuous axial scanning obtained by driving the GSM with a triangular waveform at 100Hz. Scale bar: (a-c) 10um.

To demonstrate that our focusing strategies are compatible with faster scan technologies, we replaced the GSM with a 12kHz resonant galvanometric mirror. If we scan over a 7.5 degree inclined planar mirror, the resonant scanner produces an axial line scan with a typical temporally averaged intensity profile for a sinusoidal motion. We found that we can normalize the axial intensity distribution without any synchronized laser modulation: We increased the scan range of the resonant galvo threefold, such that the approximately linear portion of the sinusoidal oscillation covered the same range as shown in Figure 3c. We placed a mask in the image space of the 4F telescope between the GSM and OBJ1, which restricted the transmitted laser beam to a smaller lateral scan range. As the lateral and axial scan ranges are coupled, this essentially allowed us to truncate the axial scan range to a smaller, more homogeneous region, at the expense of a lowered temporal duty cycle for the illumination.

**Figure 3.**
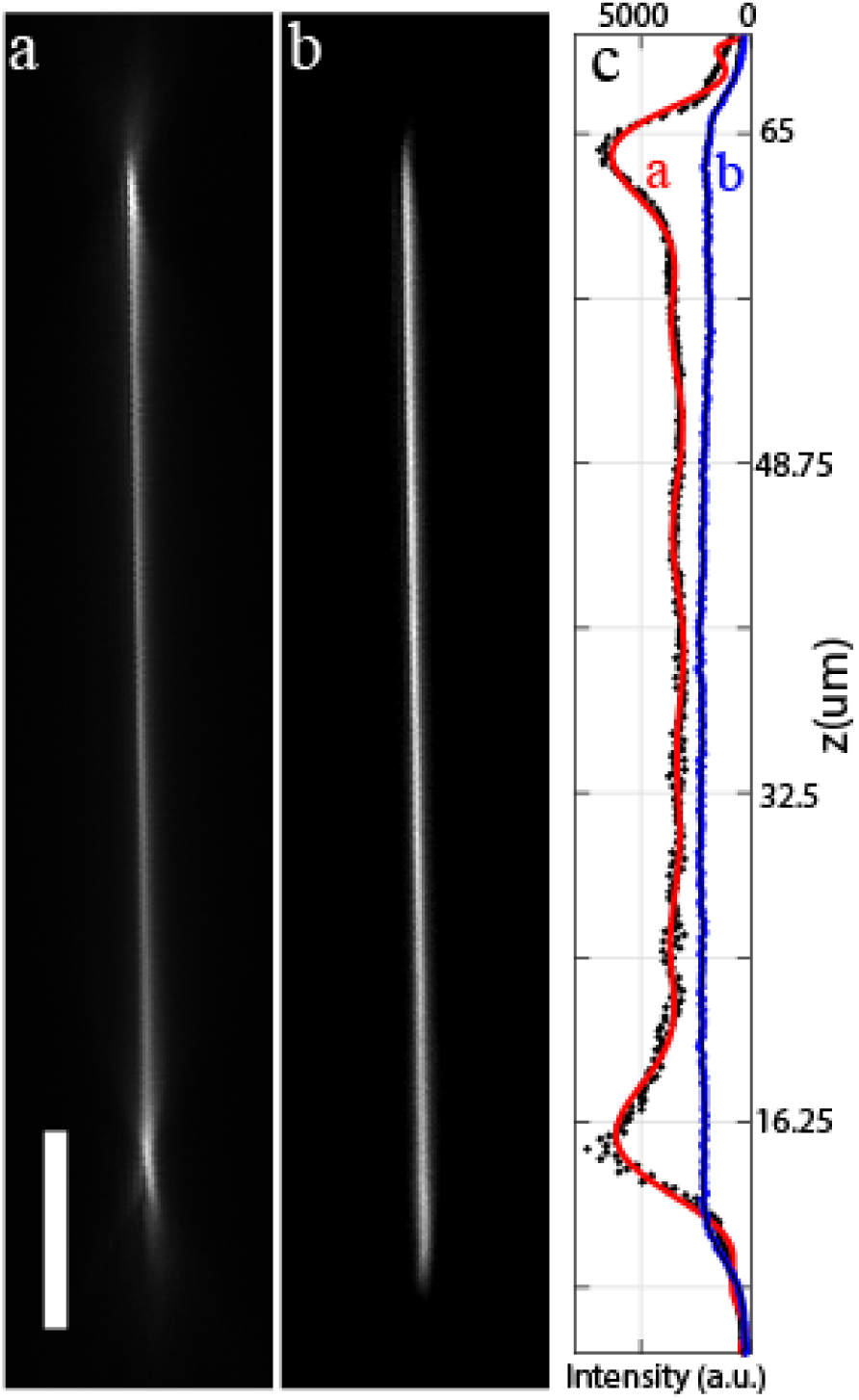
Resonant axial scanning using a tilted planar remote mirror. (a) Sinusoidal scan motion over a planar mirror inclined by 7.5 degree. (b) Same arrangement as in (a), but with a threefold increased scanning amplitude and a mask to truncate the scan range optically.

To demonstrate the usefulness of our scan technology to biological imaging, we employed our setup to perform rapid axially swept light-sheet microscopy (Figure 4a). To this end, a cylindrical lens was used to shape the input beam into a light-sheet, which was then imaged into the image space of OBJ1 (Methods). Using a planar mirror tilted by 7.5 degrees, this allowed us to scan a thin lightsheet (effective NA 0.8) over around 40 micron range. As evidenced by the point-spread function (Figure 4b), this version of ASLM can achieve slightly sub 400nm axial raw resolution, which is in accordance with previous implementations. We imaged RPE cells labeled with vimentin-GFP at both 50ms and 5ms exposure time per frame. As can be seen in figures 4c-f, the morphological detail is faithfully recovered at the tenfold faster acquisition time. To leverage this rapid ASLM system, we imaged genetically encoded multimeric nanoparticles (GEM) inside MV3 cells. Such imaging has previously only been performed two dimensionally due to restriction in volumetric acquisition speed^16^. With our system, we can acquire volumetric data (encompassing 13 individual z-planes) at a volumetric rate of 3.5Hz (XYZ image planes per second) and isotropic sub 400nm resolution over a volume spanning 128×32×2 microns and encompassing 200 timepoints. This volumetric imaging speed was sufficient to track GEMs in the perinuclear region of the cells in 3D (Figure 4 h-i).

**Figure 4.**
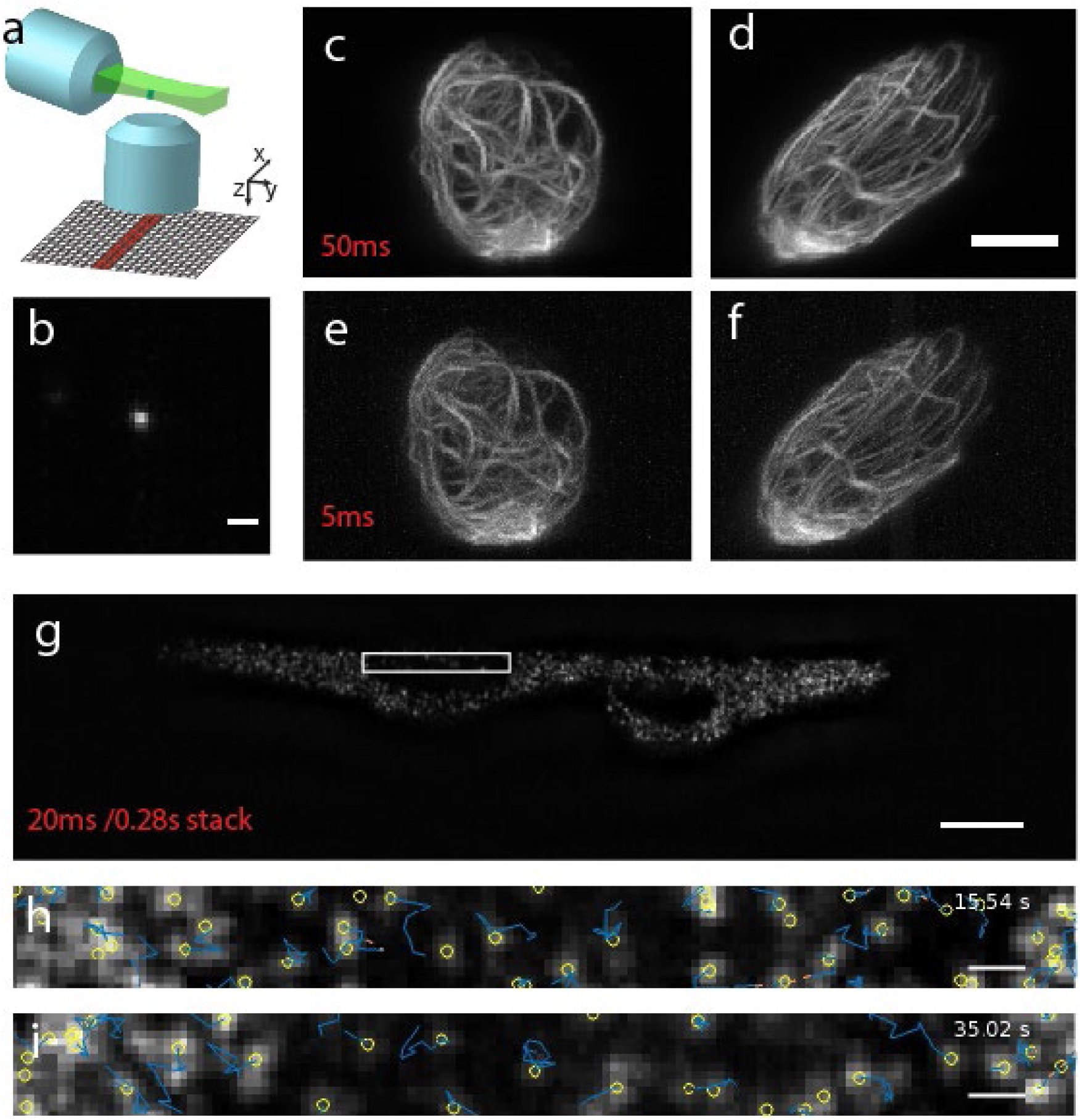
Accelerating axially swept light-sheet microscopy (ASLM). (a) schematic principle of ASLM: a thin light-sheet is scanned in its propagation direction and only the region within the beam waist (red and green bars) is read out by an sCMOS camera. (b) Point spread function in an x-z view. (c-d) RPE cell labeled with GFP vimentin, imaged with ASLM with a 50 ms integration time. x-z and y-z maximum intensity projections are shown. (e-f) The same cell imaged with ASLM at 5 ms integration time. X-z and yz maximum intensity projections are shown. (g) Genetically encoded multimeric nanoparticles (GEM) inside two MV3 cells, as imaged by ASLM at 20 ms image exposure time, which resulted in an acquisition rate of 3.57 volumes per second. (h-i) Axial y-z view of the perinuclear region at two timepoints. Yellow circles indicate detected vesicles and blue lines illustrate cumulative tracks. Scale bar (a,h,i) 1um; (d,g) 10um.

## Discussion

We present a novel approach to remote focusing that leverage lateral scan technologies to achieve rapid axial refocusing. The proposed method is compatible with even faster scan technologies than presented here, and it should be possible to reach MHz scan rates. The scan motion can be resonant or deterministic (i.e. having the ability to freely choose the scanning waveform and to perform discrete scanning steps), depending on the choice of scanning technology. This contrasts with resonantly driven ETLs, which can only provide sinusoidal waveforms. In addition, compared to the recently published reverberation microscopy, our technique can be utilized to: (a) access many more z-positions with greater flexibly, (b) is free of spherical aberrations and (c) can deliver tailored laser power to each focal plane. However, reverberation microscopy achieves much faster refocusing than presented here and can also more readily cover a larger axial scan range. Nevertheless, our method is the only one capable of performing remote focusing in an aberration-free format.

The two presented implementations, step mirror and tilted planar mirror, have different strengths and weaknesses. The step mirror allows arbitrary scan step sizes, but the field of view of the remote scanning system will put an upper limit on the number of steps that can be realized. Our concept can be extended to two scan dimensions in the remote focusing arm, allowing additional steps to be added in a second lateral dimension (suppl. Figure 7). The tilted mirror geometry allows fine axial step sizes that are in principle only limited by the angular resolving power of the lateral scan technology being used. The axial range is tied to the tilt angle and the field of view. While our scan range in this publication was limited to around 40 microns, this was mainly caused by the small FOV (∼330 microns) of OBJ1. Lenses of similar NA with much larger field numbers and corresponding tube lenses for wide fields (26mm) have become available and should increase the scan range fourfold (Supplementary note 2).

Our first practical demonstration on microscopic imaging was ASLM, which has been criticized for its slow acquisition speed (around 10Hz framerate) ^17^. Here, we demonstrate that our new scan technology allows one order of magnitude acceleration while keeping the high spatial resolving power of this emerging imaging technology. We furthermore see a major application area for our technology for multiphoton raster scanning microscopy, which is the backbone in fluorescence imaging in the neurosciences. Here, both the discrete and the continuous scan technologies presented here may find many applications to image different layers of the brain nearly simultaneously or to rapidly acquire whole volumes to measure neuronal firing patterns. For the latter, the use of resonant galvos and Lissajous scanning patterns^18^promises volumetric imaging at kHz rates. As such, we hope that this technology will find immediate and widespread applications to accelerate microscopic 3D imaging in biology and the life sciences.

## Methods

### Experimental setup

Laser light emitted from a diode pumped solid state 488 nm CW laser (Coherent Sapphire) was spatially filtered with a pinhole and expanded 9-fold. A halfwave plate was used to rotate the polarization of the beam and vary the laser power. In the remote focusing arm, we used a 60× 0.7 NA air objective (Nikon), two f=200 mm telecentric tube lenses (AC508-200-A, ThorLabs), a quarter-wave-plate QWP, (Newport) and a Galvo scanner (CRS12kHz, Cambridge Techologies). As this air objective required a coverslip, a 170-micrometer thick coverslip was glued to the front surface of the lens. The illumination arm consisted of a 40X NA 0.8 water dipping objective (Nikon), and two achromatic doublets (AC508-150-A and AC508-300-A, ThorLabs)). A custom-made sample chamber was used into which the water dipping lens was immersed. The relay lenses were chosen such that the overall demagnification from the remote mirror to the water chamber corresponds to 1.333. A traditional light-sheet detection arm was oriented at 90 degrees (see also Figure 1) and consisted of a 40X NA 0.8 water dipping objective (Nikon), a f=200 mm telecentric tube lenses and an Orca Flash 4 camera. For fluorescence detection, a long pass filter (Semrock) was used to block the laser light, and the sample was scanned through the ASLM illumination beam with a piezo stage (PIHera, P-621.1CD, Physik).To evaluate the beam in transmission, another detection arm was built with neutral density filters to attenuate the laser. Objective piezo (PIFOC, Physik Instrumente) was used to acquire 3D stacks of the laser spots using z-stepping of the objective (i.e. the tertiary objective corresponding to the imaging arm). For the fluorescence detection arm, no provisions for z-stepping of the objectives were made.

### Focal spot measurements

To measure the optical quality of our remote focusing technique, we acquired transmission datasets with the tertiary imaging system shown in Figure 1. To this end, a tertiary imaging objective was carefully aligned along the optical axis of OBJ2 and an image of the laser spot was formed on the camera. To acquire 3D data, we stepped the tertiary objective with the objective piezo. The step size was chosen to 160 nm, which resulted in isotropic voxel sizes, and typically, a volume with a z-range of 100 micron was acquired, which encompassed the full refocus range used in this work. This procedure was repeated for each focal position when refocusing was performed in discrete steps (i.e. when the regular galvanometric mirror was used in discrete steps). When the galvo was driven by a triangular waveform or the resonant scanner was used, similarly, a 3D transmission stack was acquired. Since the resonant scanning occured much faster than the acquisition of the 3D stack, a time averaged intensity distribution of the rapidly moving focus is acquired.

### Mammalian cell culture and labeling

The hTERT immortalized human Retinal Pigment Epithelial cells (hTERT-RPE-1) were obtained from ATCC (Manassas, VA) and grown in DMEM/F12, 1:1,(GIBCO, Life Technologies) medium supplemented with 10% fetal bovine serum(GIBCO), penicillin/streptomycin(GIBCO) and 0.01mg/ml hygromycin-B (Sigma, St. Louis MO). Cells were tagged with mEmerald-vimentin using TALEN based genome editing approach which was previously described^19^. For imaging, cells were placed into collagen as previously described^14^. MV3 cells were obtained from Peter Friedl (MD Anderson Cancer Center, Houston TX). MV3 cells were cultured in DMEM (Gibco) supplemented with 10% fetal bovine serum (FBS; ThermoFisher) in 37°C and 5% CO2. Cells were infected to express genetically encoded multimeric nanoparticles (GEMs) using lentiviral construct from Addgene (Plasmid #116934^16^). Cells were FACS sorted to purify population of cells expressing GEMs.

## Acknowledgment

This research was funded by grants from the Cancer Prevention Research Institute of Texas (RR160057 to R.F.) and the National Institutes of Health (F32GM117793 to K.M.D., R33CA235254 to R.F.).

## Supplementary notes

### Note 1, Simulation of the remote focusing setup using Zemax

This supplementary note describes the results of a simulation of the remote focusing setup with a tilted mirror using Zemax Optic Studio 18.4.1. The simulation comprised of a beam splitter, a galvo-scanning mirror, a remote mirror, four achromatic doublets (Zemax model of AC508-200-A, Thorlabs) for two 4f systems, and two water immersion 40x NA 0.8 objective models which were kindly provided by Ryan McGorty [1] (Fig. S3). It has to be noted that in our experimental setup, we use an air objective as the remote objective, which however was not available to us as a model for simulations. To find the right distances (thicknesses) of the optical surfaces in the simulation, we applied optimization functions available in Zemax such as “Quick Adjust” and the “Merit Function Editor” and manually changed parameters using the Slider adjustment in the optimization tab of Zemax.

In the simulation, an incoming collimated beam with defined aperture (Fig. S4) was reflected 90° degrees by a mirror onto a galvo-scanning mirror tilted by 45°□ 1.5° degrees corresponding to □ 3° optical scan angle. The galvo-scanning mirror was conjugated to the back focal plane of a remote focusing objective (40x, NA 0.8) with a telescope of two achromatic doublets of two inch diameter with f=200 mm (AC508-200-A, Thorlabs). The remote focusing objective focused the beam on a remote mirror which was tilted by 2.5°, 5° or 7.5° degrees. To ensure that the incoming beam was reflected onto the same beam path, we optimized the chief ray to be reflected exactly onto the incoming beam by adding an off-axis shift *d* to the objective. The reflected beam travelled back through the above system (objective and 4f system) and was descanned by the galvo-mirror. Then, the galvo-mirror was conjugated with another 4f-system to the back focal plane of the imaging objective. Scanning the beam in the remote focus objective over the tilted mirror, therefore scanned the beam position along the imaging axis. For each mirror tilt (2.5°, 5°, 7.5° degrees), we determined the final focus position manually for 7 galvo-mirror scan positions (−2°,-1.5°,-1°, 0°, 1°, 1.5°, 2° scan mirror tilt). To the obtained values, we fitted a 4^th^ degree polynomial (Fig. 5A) to determine for each mirror tilt the corresponding focus position within the scan range of the imaging objective. This enabled generating automatic measurements of the PSF and their width as well as the Strehl ratio along the scan direction using Zemax’s inherent ZPL programming language (Fig. S4, S5, S6). Simulation results showed that the PSF in y-direction of the laser foci along the scanning direction in the imaging space was skewed (Fig. S5F). By manually adjusting the off-axis shift *d* of the objective, this tilt could be straightened (Fig. S6F) at the cost of a small decrease of resolution (Fig S6).

### Note 2: Axial scan range using a tilted mirror

The change of focus dz that can be achieved with a tilted mirror is given:

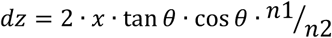

Where θ is the tilt angle of the mirror, n1 is the refractive index of the immersion media of the remote objective and n2 is the refractive index of the immersion media of Objective 2.

In our setup, we used a 60X objective and the tube lenses limited us to a FOV of 20m, which equals to a scan range over the remote mirror of ∼333microns. According to the above formula, this corresponds to a maximum axial scan range of 65 microns with a tilt angle of 7.5 degrees. In practice, we noted that at the end of the scan range, field curvature imposed a notable tilt onto the final focus and a more practical linear range was around ∼40 microns.

The tilt angle itself is limited by the remote objective: in order to use the full NA of objective 2 for focusing, the remote objective needs to possess a larger angular aperture than Objective 2. In our experimental example, the half opening angle of the NA 0.7 air objective is 44.4 degrees and the half angle of Objective 2 (NA 0.8, water) is 36.9 degrees, which allows us to tilt the beam by ∼7.5 degrees.

There are higher NA air objectives (as high as NA 0.95) and hence potentially larger tilt angles could be used to increase the axial scan range. However, we believe a better strategy is to keep the tilt angle small and instead increase the field of view of the remote objective.

Indeed, recently a 20X NA 0.8 air objectives and matching tube lenses with a field of view of 26.5mm have been developed (e.g. Olympus UPLXAPO20X and SWTLU-C). Building a remote focusing arm with these components and using a scan range of 1.325 mm and a tilt of 7.5 degrees would enable an axial scan range of 260 microns.

### Note 3: Geometrical considerations for the step sizes of the mirror

Here we analyze the relationship of the step size of the mirror in relation to its height and the half opening angle of the laser focus, using geometrical optics as an approximation In supplementary figure 7 it can be seen that the minimal width dx for the n-th step with height dz and the half opening angle α scales in the in the following:

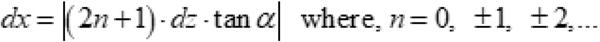

Here n=0 correspond to the nominal focal plane.

This makes it obvious that increasingly wider steps have to be used with increasing defocus. To give the reader an idea how many steps can be realistically fit into a field of view, we provide a few examples, where we numerically filled increasingly wider steps into a fixed interval.

For our setup, a half opening angle of α=36 degrees and a field of view / scan range of 330 micron is assumed for our remote focusing objective. To achieve 1 micron z-steps in the primary objective, a mirror step size dz of 0.6666 microns is required. For such a step, using the above relationship for the width of the step sizes, we found that one can fit 9 steps.

Using recently developed large Field number objectives and tube lenses (UPLXAPO20X and SWTLU-C), a FOV of 1.325 is possible, which would result in 13 steps.

## Supplementary Figures

**Supplementary Figure 1.**
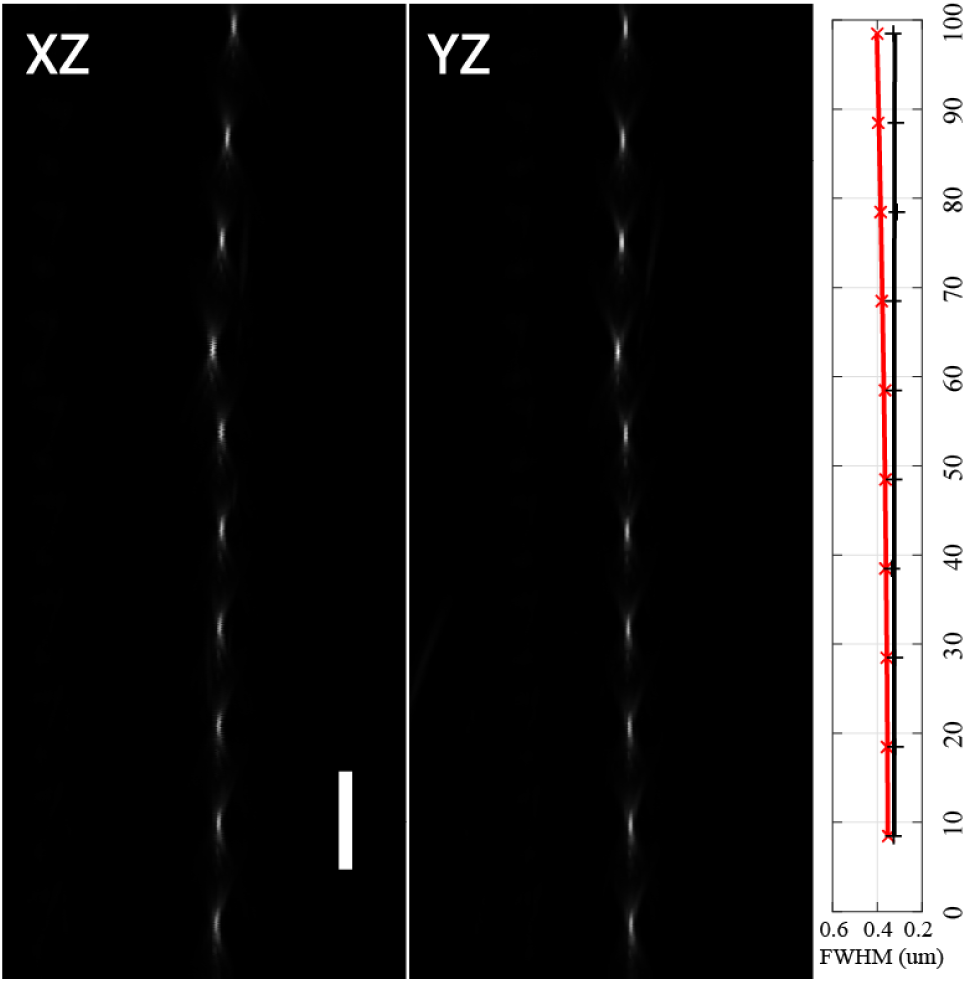
XZ and YZ view of remote focusing using a planar mirror at zero degree tilt, which was mechanically stepped in the axial direction. This method is expected to be aberration free and serves as a reference for focus quality. Scale bar: (a-d) 10um.

**Supplementary Figure 2.**
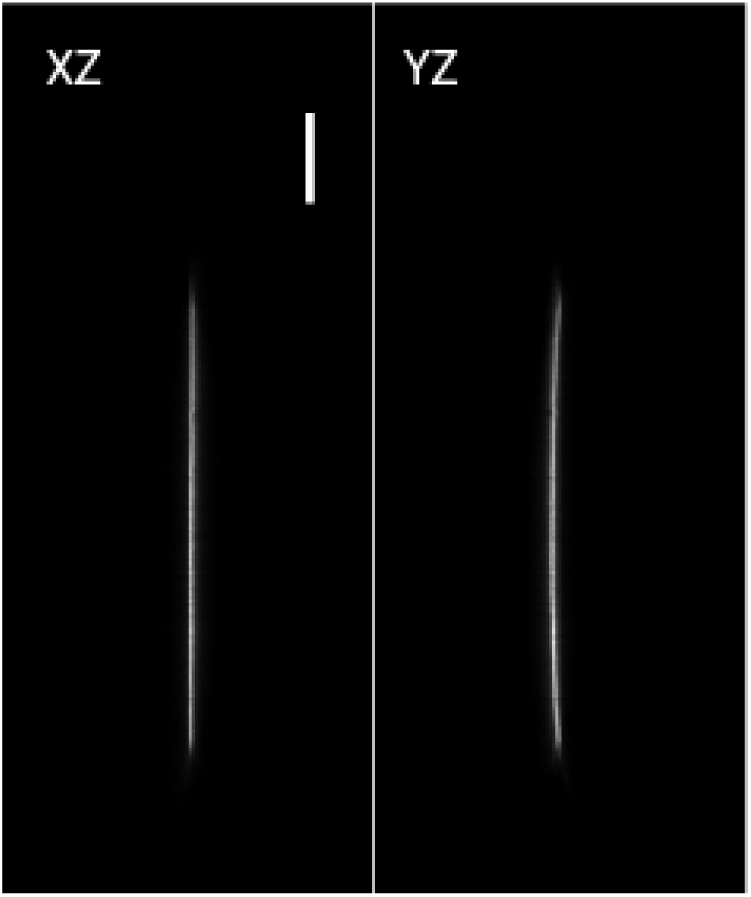
XZ and YZ view of continuous axial scanning obtained using a mirror tilt of 7.5 degree by driving the GSM with a triangular waveform at 100Hz. The YZ view shows the bent in the axial scan when the edges of the field of view of the remote scanning system are reached. Scale bar: (a-d) 10um.

**Supplementary Figure 3.**
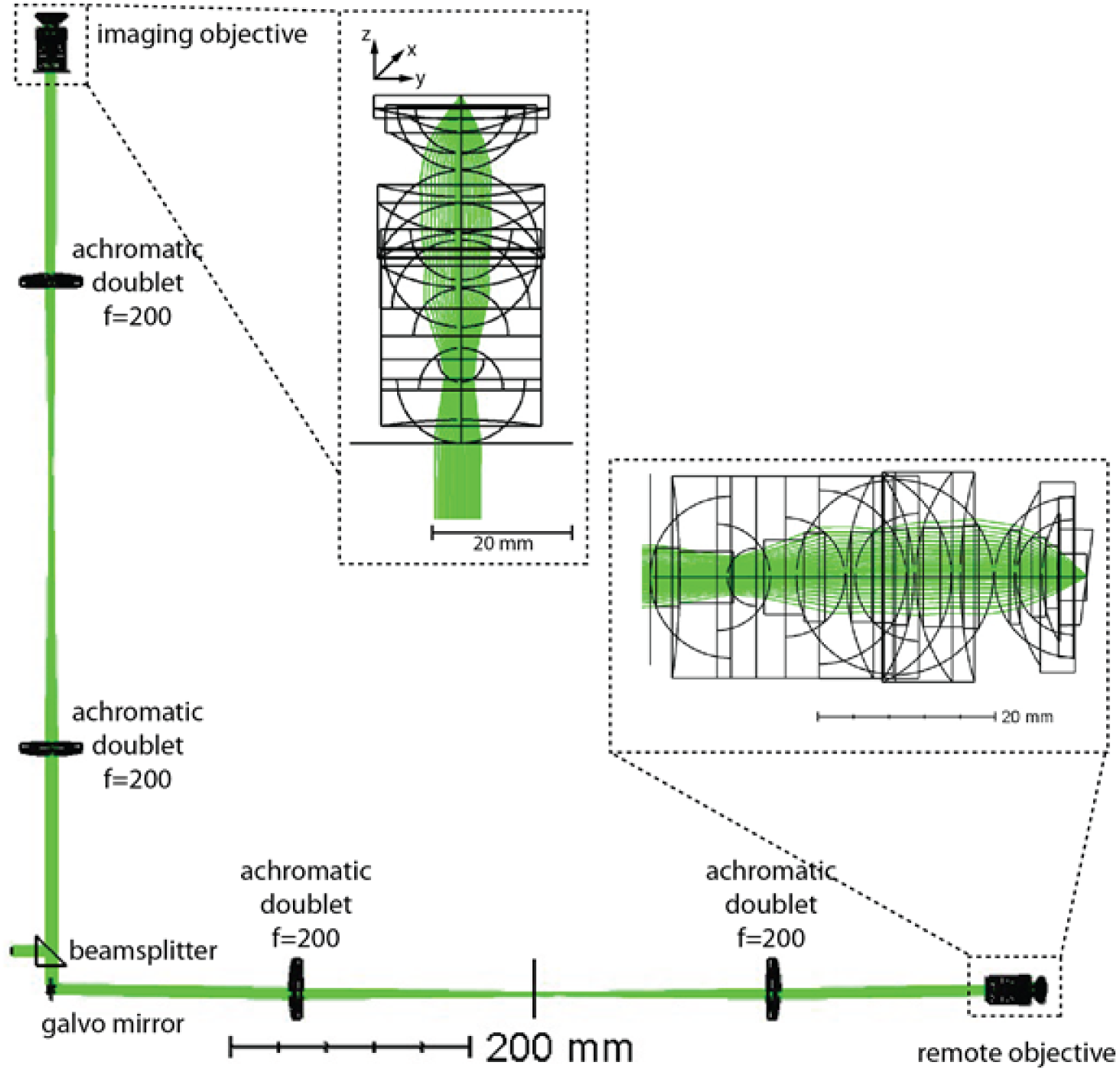
Schematic overview of the simulation. Renderings from Zemax 3D viewer as overview about the simulation (scale bar 200 mm). Inlets highlight the remote objective and the imaging objective (scale bar 20 mm). Here, a galvo-mirror scan angle of 1,152° degrees is displayed. The incoming beam was set at an aperture of 6.43 mm, and the mirror tilt was set to 7.5° degrees.

**Supplementary Figure 4.**
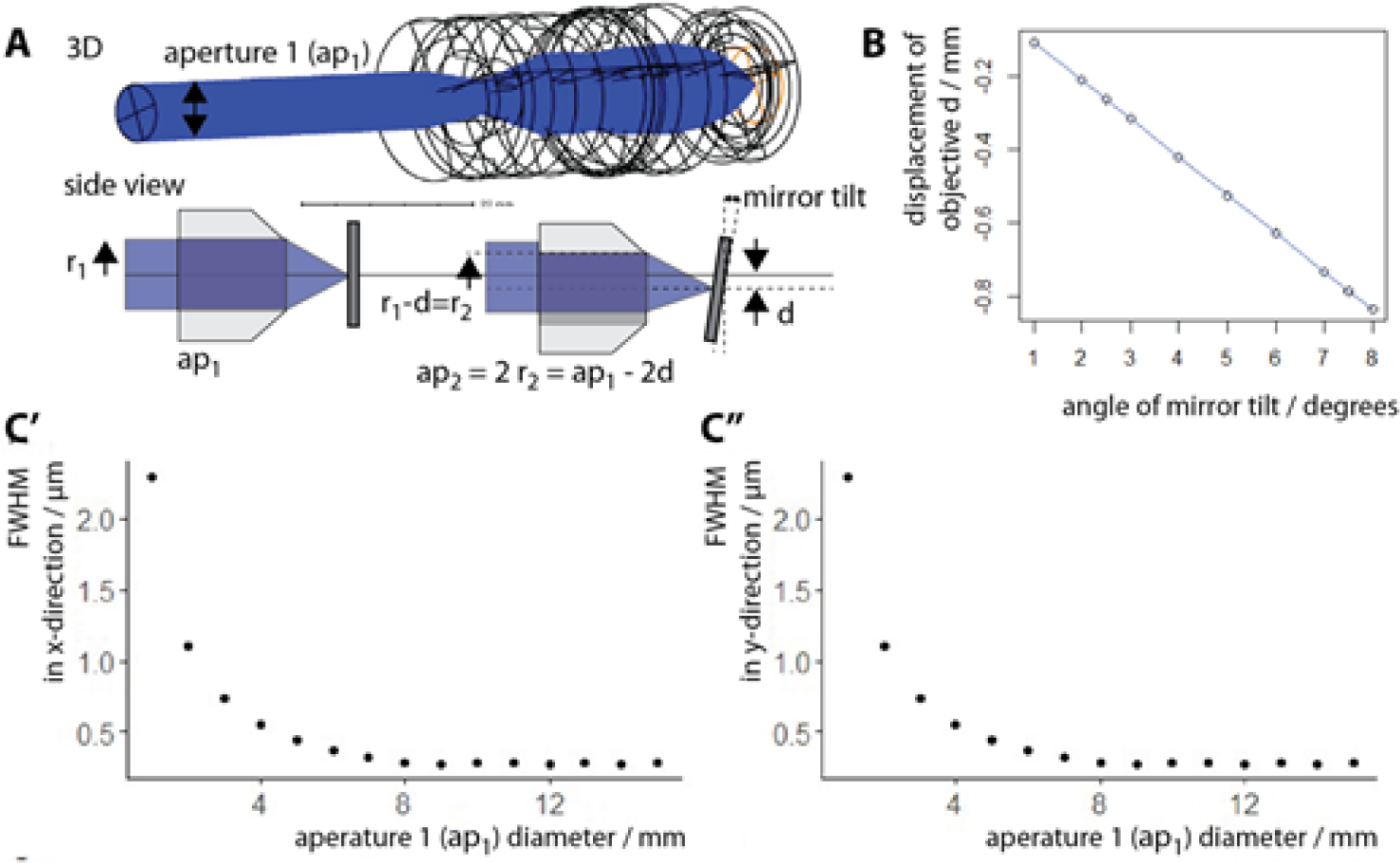
Determination of aperture of incoming beam. **(A)** Schematic drawing of dependence of the maximal available aperture of the imaging system in relation to the mirror tilt (*top*: aperture ap_1_ in case of straight mirror, *bottom*: schematic comparison of straight vs tilted mirror). To ensure that the outcoming beam was reflected on the same optical path as the incoming beam, the objective was displaced from the central optical axis. Objective displacement (d), however, reduced the maximal available aperture (ap_2_ with radius r_2_) in relation to the aperture of the system with a straight mirror (ap_1_ with radius r_1_) according to the relation ap_1_ – 2d. Therefore, to determine the available aperture for a given mirror tilt, the required displacement (d) and the unperturbed aperture (ap_1_) had to be determined for the given objective. **(B)** We measured the required displacement of the objective d in relation to the mirror tilt by overlapping the chief ray of the outcoming beam (after the mirror) onto the chief ray of the incoming beam (black dots). The dependence was linear (blue line, R-squared =1). **(C’, C”)** To determine the maximal available aperture of the objective, we determined the FWHM of the PSF in x-direction (C’) and y-direction (C”). After an aperture of 8 mm, the FWHM did not decrease anymore. Therefore, we set ap_1_= 8 mm.

**Supplementary Figure 5.**
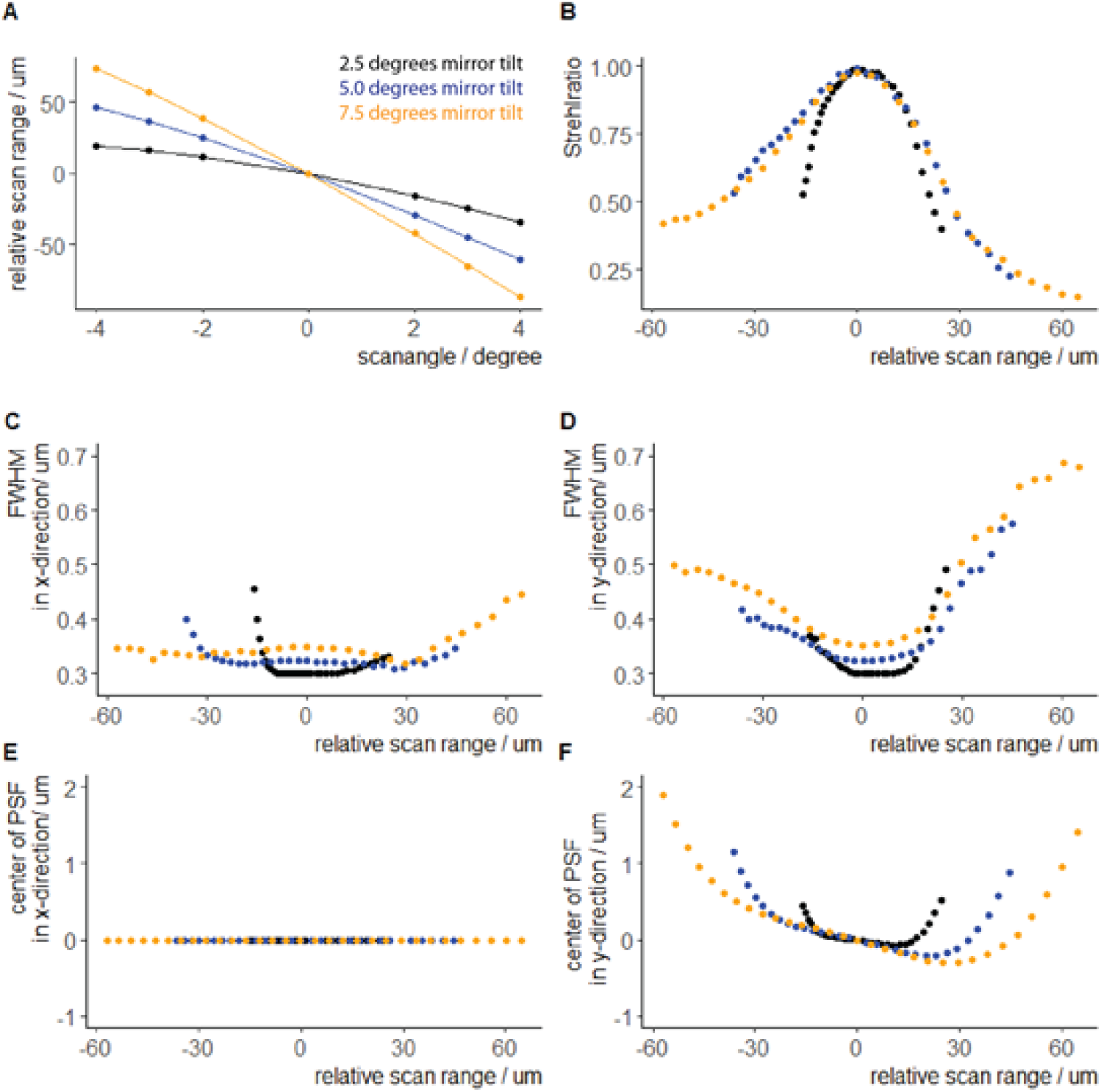
Simulation PSF measurements over scan range. We measured the properties of the Huygens PSF of the simulated imaging system at the focus of the beam in the imaging chamber at 2.5° (black), 5.0° (blue) and 7.5° (orange) degrees mirror tilt of the remote mirror over a range of □ 3° optical scan angle of the galvo-mirror. In this setup, the chief ray of the incoming beam was matched onto the chief ray of the reflected beam at the mirror. **(A)** Dependence of the relative scan range on the galvo-mirror scan angle. We measured the values at-2°,-1.5°,-1°, 0°, 1°, 1.5°, 2° scan mirror tilt (points) and fitted a 4^th^ degree polynomial to the values (line). **(B)** Changes of the Strehlratio over the relative scan range. **(C)** Full width half maximum (FWHM) measured at the center of the PSF in x-direction. **(D)** Full width half maximum (FWHM) measured at the center of the PSF in y-direction. **(E)** Shift in the center of the Huygens PSF in x-direction over the simulated scan range. **(F)** Shift in the center of the Huygens PSF in y-direction over the simulated scan range.

**Supplementary Figure 6.**
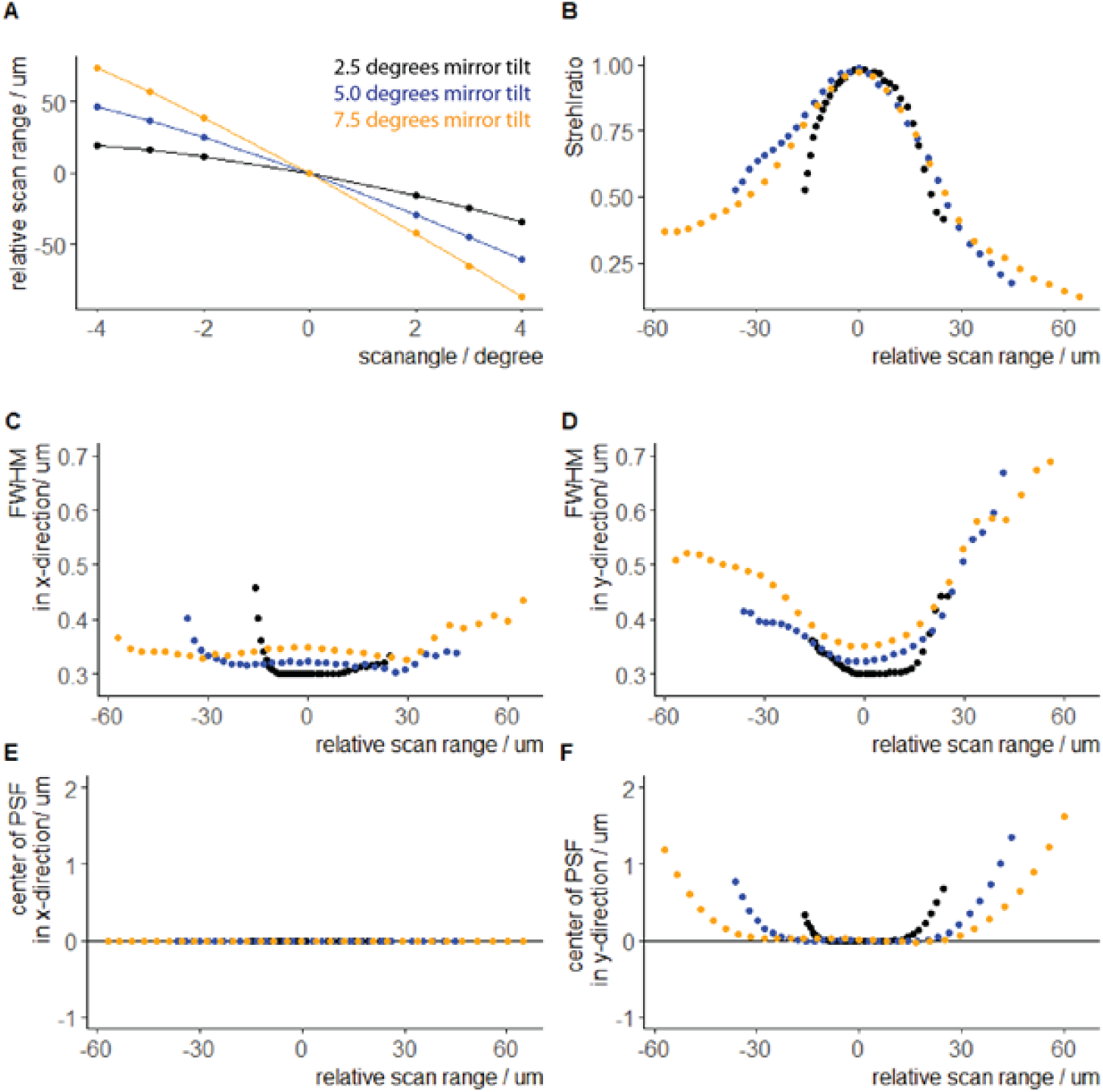
Simulation PSF measurements over scan range in optimized setup. We measured the properties of the Huygens PSF of the simulated imaging system at the focus of the beam in the imaging chamber at 2.5° (black), 5.0° (blue) and 7.5° (orange) degrees mirror tilt of the remote mirror over a range of □ 3° optical scan angle of the galvo-mirror. In this setup, the objective displacement was optimized to place the focus of the PSF onto the geometric center (x-shift = 0 mm, y-shift=0 mm). **(A)** Dependence of the relative scan range on the galvo-mirror scan angle. We measured the values at-2°,-1.5°,-1°, 0°, 1°, 1.5°, 2° scan mirror tilt (points) and fitted a 4^th^ degree polynomial to the values (line). **(B)** Changes of the Strehlratio over the relative scan range. **(C)** Full width half maximum (FWHM) measured at the center of the PSF in x-direction. **(D)** Full width half maximum (FWHM) measured at the center of the PSF in y-direction. **(E)** Shift in the center of the Huygens PSF in x-direction over the simulated scan range. **(F)** Shift in the center of the Huygens PSF in y-direction over the simulated scan range.

**Supplementary Figure 7.**
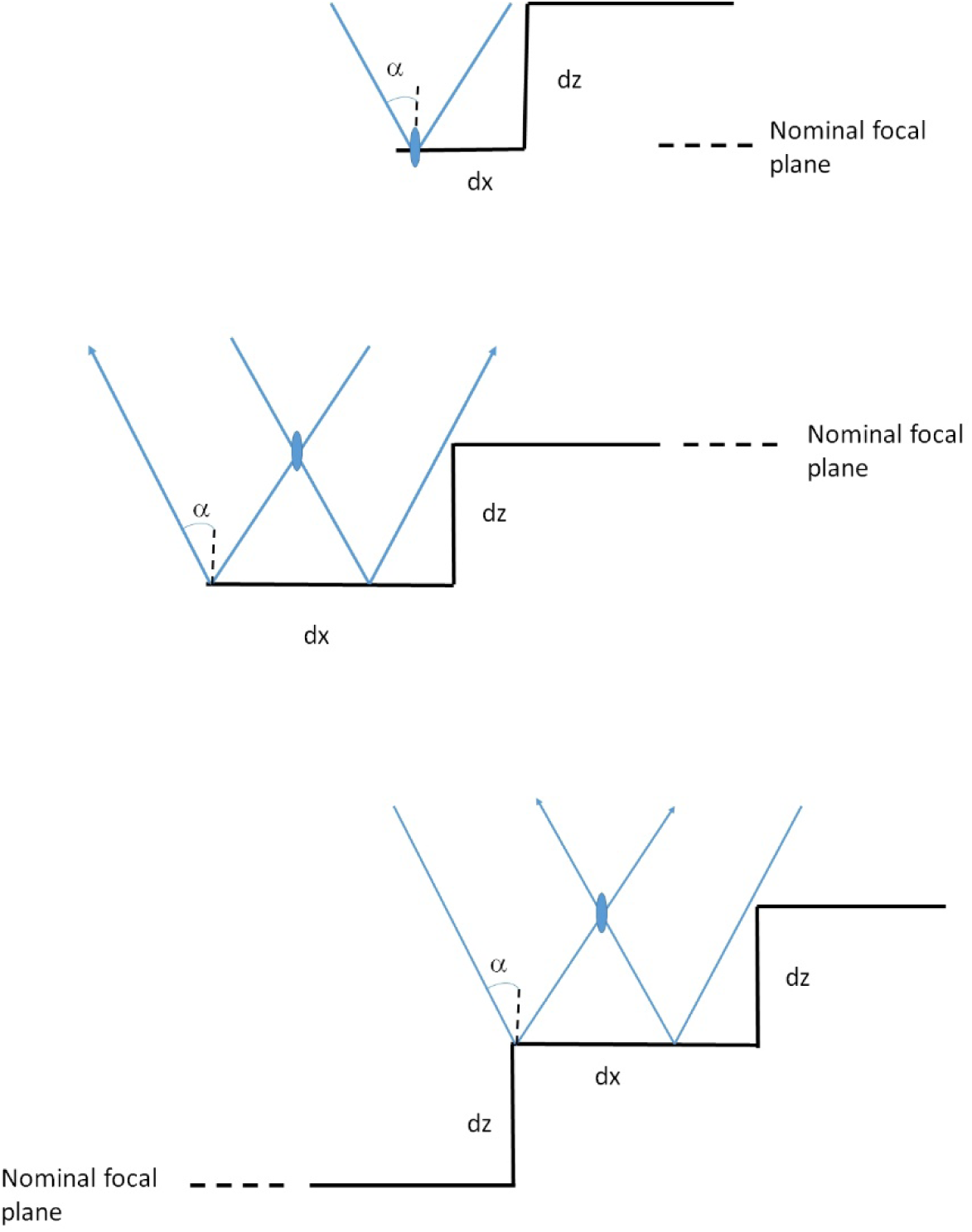
Geometrical considerations for the width (dx) of the mirror steps in relation to the half opening angle α and the step height dz. Top graphic shows the scenario where the step is in the nominal focal plane. Middle graphic shows a step that is below the nominal focal plane and bottom graphic shows the scenario where the step is above the nominal plane. Blue lines depict the marginal ray of the laser focus and arrows indicate the propagation direction of the light.

**Supplementary Figure 7.**
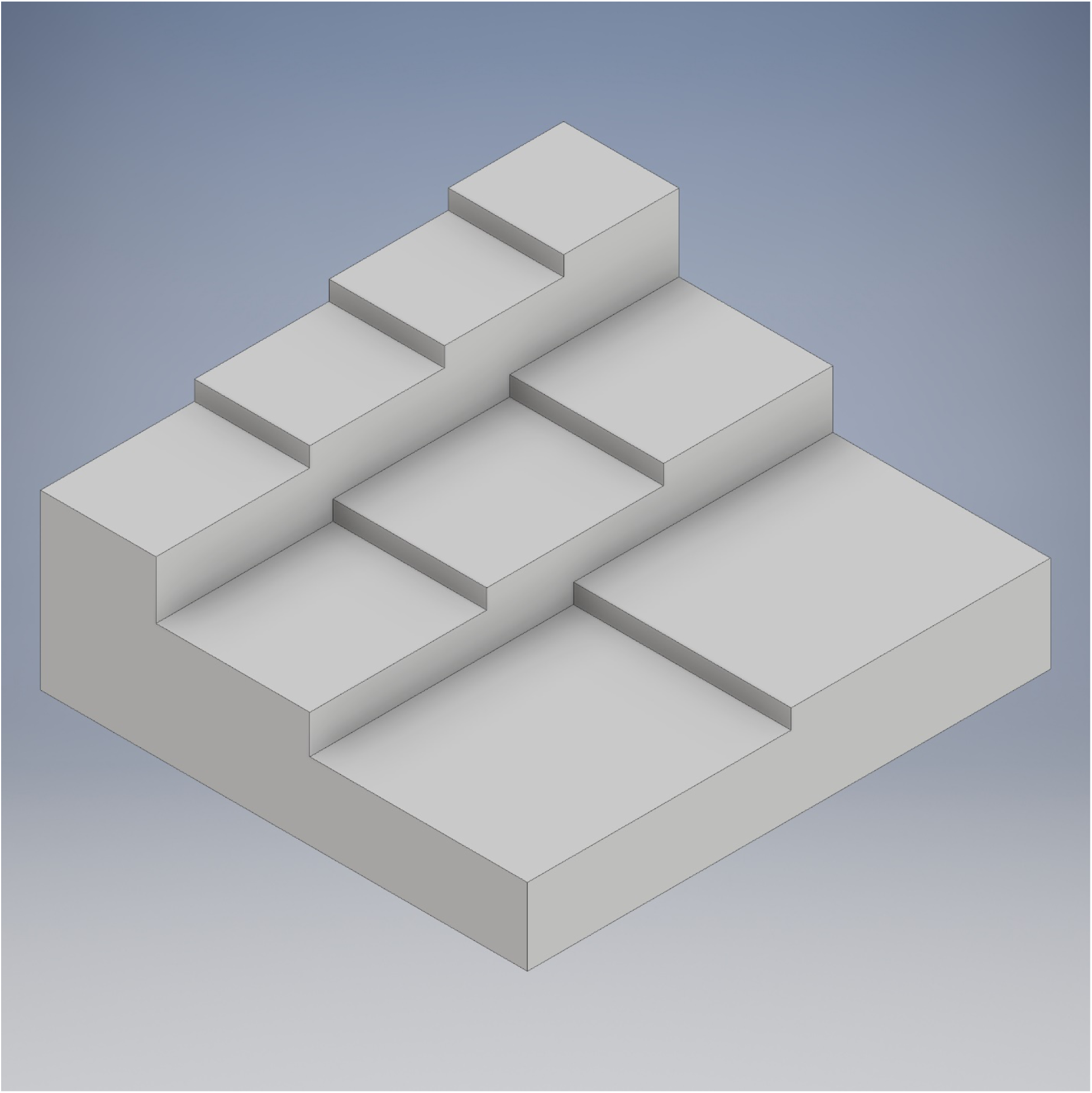
3D rendering of a two-dimensional stair case mirror with 9 steps. Steps widen with increasing axial distance from nominal focal plane. A two dimensional raster scan over this mirror could address 9 different focal planes.

